# The Ulcerative Colitis-Associated Gene NXPE1 Catalyzes Glycan Modifications on Colonic Mucin

**DOI:** 10.1101/2024.12.24.630288

**Authors:** Ranad Humeidi, Noriko Rapley, Xiebin Gu, Joon Soo An, Ashwin N. Ananthakrishnan, Elizabeth A. Creasey, Mark J. Daly, Stuart L. Schreiber, Daniel B. Graham, Mohammad R. Seyedsayamdost, Ramnik J. Xavier

## Abstract

Colonic mucus forms a first line of defense against bacterial invasion while providing nutrition to support coinhabiting microbes in the gut. Mucus is composed of polymeric networks of mucin proteins, which are heavily modified post-translationally. The full compendium of enzymes responsible for these modifications and their roles in health and disease remain incompletely understood. Herein we determine the biochemical function of NXPE1, a gene implicated in ulcerative colitis (UC), and demonstrate that it encodes an acetyltransferase that modifies mucin glycans. Specifically, NXPE1 utilizes acetyl-CoA to regioselectively modify the mucus sialic acid, 5-*N*-acetylneuraminic acid (Neu5Ac), at the 9-OH group to generate 9-*O*-acetylated Neu5Ac (Neu5,9Ac_2_). We further demonstrate that colonic organoids derived from donors harboring the missense variant NXPE1 G353R, which is protective against UC, exhibit severely impaired acetylation of Neu5Ac on mucins. Together, our findings support a model in which NXPE1 masks the alcohols of mucus sialoglycans via acetylation, which is important for modulating mucus barrier properties that limit interactions with commensal microbes.

## INTRODUCTION

Inflammatory bowel disease (IBD) is a group of disorders comprised of Crohn’s disease (CD) and ulcerative colitis (UC), which involve chronic inflammation of the digestive tract. In 2020, there were ∼2.5 million IBD patients in the United States, a number that is estimated to rise to ∼3.5 million by 2030 (1). IBD can be debilitating and can lead to life-threatening complications. Studies have linked IBD to specific environmental conditions, genetic risk factors, and immune dysregulation, which suggests a complex pathophysiologic process (2–4). Genome-wide association studies (GWAS) have been particularly successful in identifying genetic loci associated with CD and UC, some of which can be fine-mapped to implicate causal genes and variants (5–9). One such gene, *NXPE1* (formerly FAM55A), encodes for a previously uncharacterized protein that has a missense variant protective against UC through yet unknown mechanisms (6). Characterization of these variants, therefore, can provide insights into IBD as well as new molecular targets and possible treatment options (10).

Based on its protein sequence and homology to known enzymes, NXPE1 is classified as a putative member of the GDSL/SGNH esterase superfamily and, within it, the PC-esterase family. Members of these families are serine hydrolases, one of the largest functional enzyme classes in mammals comprising 1-2% of the total proteome (11). Serine hydrolases feature a conserved Gly-Asp-Ser-Leu motif and a Ser-Gly-Asn-His active site, after which the superfamily is named, and are capable of catalyzing multiple reactions in diverse metabolic processes (12). PC-esterases have been purified and characterized from plants and bacteria, where they play roles in mediating surface glycan acylation levels, a feature key to defining chemical properties of glycans such as tensile strength, metabolite recycling, hydrophobicity, and susceptibility to degradation (11, 13, 14). In animals however, only one family member has been characterized thus far, CASD1, an acetyltransferase that specifically catalyzes the modification of sialic acids, which are essential carbohydrates that cap glycoproteins and glycolipids (15). This process is linked to cell recognition and, importantly, pathogen resistance mechanisms. Other human proteins that may be involved in this process have yet to be identified (16, 17).

The primary barrier between epithelial cells of the intestinal lining and the microbial flora in the gut is the mucosal layer, which consists of highly glycosylated proteins (18–20). Defects in this layer and the subsequent loss of barrier integrity result in dysregulation of intestinal homeostasis and are known to trigger an inflammatory response (21–24). Intestinal mucus is primarily composed of networks of mucin glycoproteins containing numerous oxygen-linked glycans that may be further modified (25–27). Due to the dynamic nature of gut mucus, these glycoproteins are heavily and reversibly modified post-translationally (23, 28). However, the enzymes responsible for these modifications are incompletely described (29, 30). Given the putative enzymatic function of NXPE1 and its association with UC, we hypothesized that NXPE1 may regulate the production and maintenance of the mucosal barrier, thus playing a crucial role in controlling the gut immune response. We, therefore, initiated a multi-pronged study to better understand its role in UC.

Here, we report that NXPE1 is an *O*-acetyltransferase of mucosal sialic acids and is involved in the production and maintenance of the mucosal barrier. NXPE1, is thus an important factor in mediating the gut immune response. These findings are validated by organoid studies, which revealed significant impairment of sialic acid acetylation in organoids carrying the UC-protective variant or in NXPE1 knockouts. Our results thus provide a mechanistic basis for the protective effect of NXPE1 variants. Moreover, they illustrate the power of deorphanizing enzymes encoded in the human genome to gain mechanistic insights into human biology and disease.

## RESULTS

### NXPE1 is highly expressed in colonic epithelial cells

To begin, we sought to determine the expression levels and localization of NXPE1 in epithelial cells and lineages thereof, which are the main source of mucus in the intestinal lining (31). Analysis of previously available, spatially defined single-cell RNA sequencing (scRNA-seq) data from patient-derived biopsies (32) revealed the highest expression of NXPE1 in goblet cells (GCs), transit amplifying (TA) cells, intestinal epithelial stem cells, and absorptive enterocytes (**Figure 1A**). The expression of other NXPE paralogs, such as NXPE2 and NXPE4, tracked with that of NXPE1 (**Figure 1A**).

**Figure 1.**
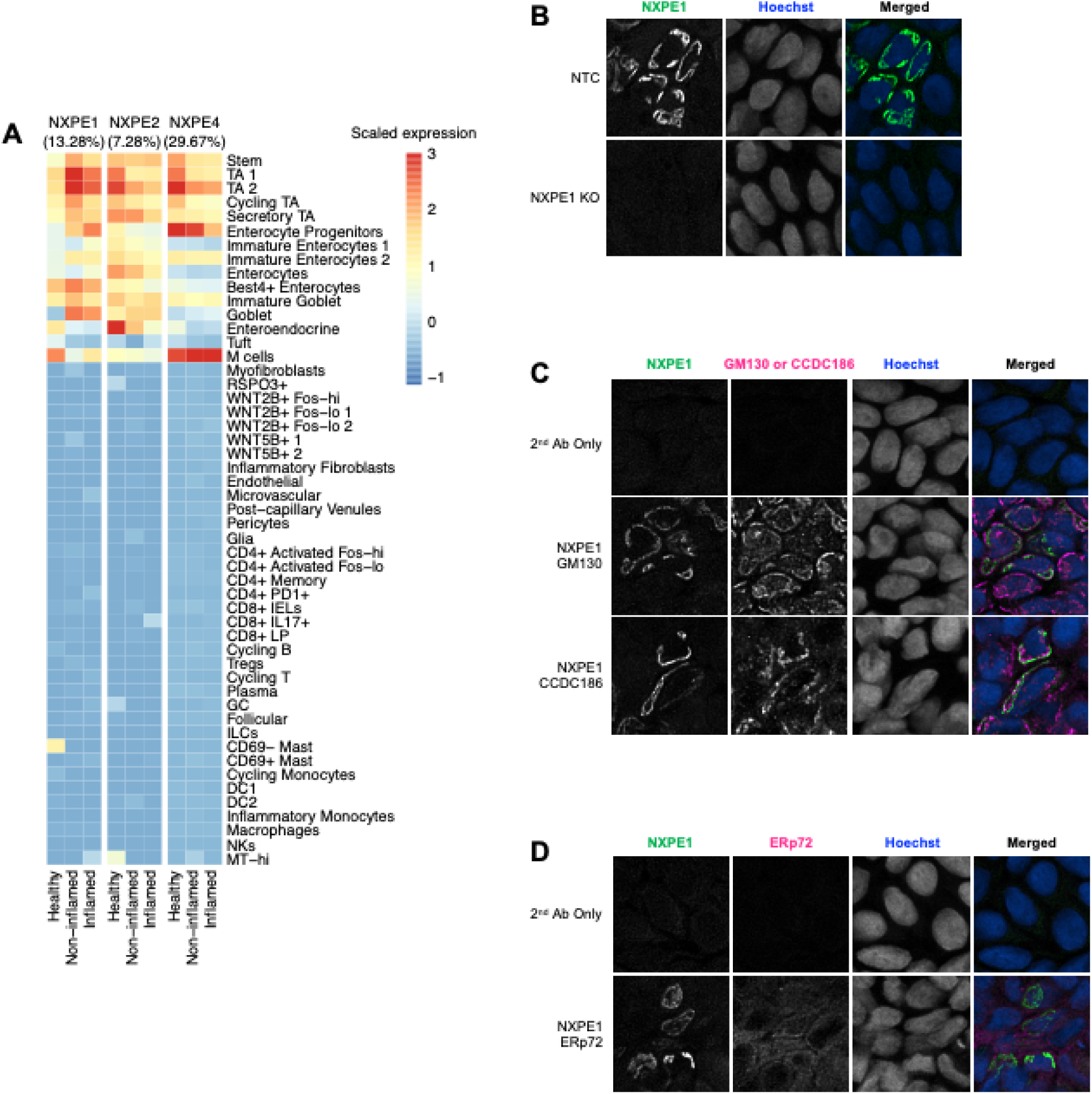
NXPE1 is expressed in intestinal epithelial cells within the Golgi apparatus. (**A**) Heatmap showing expression levels of NXPE1, NXPE2, and NXPE4 from human colon biopsies (32). Percentages in parentheses indicate the mean percent of epithelial cells in which the transcript is detected. (**B**) Detection of NXPE1 with monoclonal antibodies in human colonic organoids grown in monolayers. Staining conducted for DNA (Hoechst) and NXPE1 is shown in organoids after CRISPR/Cas9 modification with nontargeting control (NTC) or single guide RNA (sgRNA) targeting NXPE1 (KO). (**C**) NXPE1 co-localization with Golgi markers GM130 and CCDC186 in human colonic organoids. (**D**) NXPE1 co-localization stain with the ER marker ERp72 in human colonic organoids.

As further evidence, we examined the localization of NXPE1 in situ by staining GC-polarized intestinal organoids from human donor-derived biopsies. Using CRISPR/Cas9, NXPE1^−/−^ organoids were generated as a control. We observed increased expression of NXPE1 in GC-differentiated organoids compared to undifferentiated stem cells. To analyze the subcellular localization of NXPE1, we performed immunofluorescence staining and imaging studies. First, we generated a monoclonal NXPE1 antibody for immunofluorescence imaging and showed that it specifically detected NXPE1 in organoids, while the signal was ablated after CRISPR-mediated knock-out of NXPE1 (**Figure 1B**). Next, we focused on locating cellular compartments known to control protein glycosylation and co-stained with the ER marker ERp72 and the Golgi markers GM130 and CCDC186. Accordingly, we found that NXPE1 localizes to the Golgi apparatus rather than the ER (**Figure 1C and D**). Together, these results show that NXPE1 is highly expressed in colonic epithelial cells and localizes to the Golgi compartment.

### NXPE1 is an *O*-acetyltransferase of the serine hydrolase family

Based on its protein sequence and homology to known enzymes, NXPE1 has been classified as a serine hydrolase and a member of the GDSL/SGNH esterase superfamily (**Figure 2A**) (33). The UC-protective variant carries a modification in this sequence, converting the essential glycine in the GDSL motif (G353) to arginine (G353R). Moreover, sequence analysis shows that serine-355 is conserved and likely acts as the active site covalent catalyst. To determine whether NXPE1 is indeed a serine hydrolase, we used an established activity-based protein profiling (ABPP) assay, wherein the putative active site serine is specifically and covalently labeled with a fluorophosphonate-based probe that is then detected through a biotin tag (**Figure 2B**) (34). A control experiment with a validated serine hydrolase, the bovine coronavirus esterase (BCovHe), showed expected labeling and the double-band pattern previously described, representing the monomeric and dimeric forms of the protein (35–37). Likewise, experiments with NXPE1, but not the S355A variant, revealed labeling with the ABPP probe (**Figure 2B**). These studies were not possible with the NXPE1 G353R variant, as it was insoluble in both mammalian and bacterial expression systems (**Figure 2C**). In fact, other substitutions at G353, such as a tryptophan, lysine, asparagine, and even alanine, resulted in insoluble protein. These results point to the crucial role of G353 in maintaining an intact and folded NXPE1 structure.

**Figure 2.**
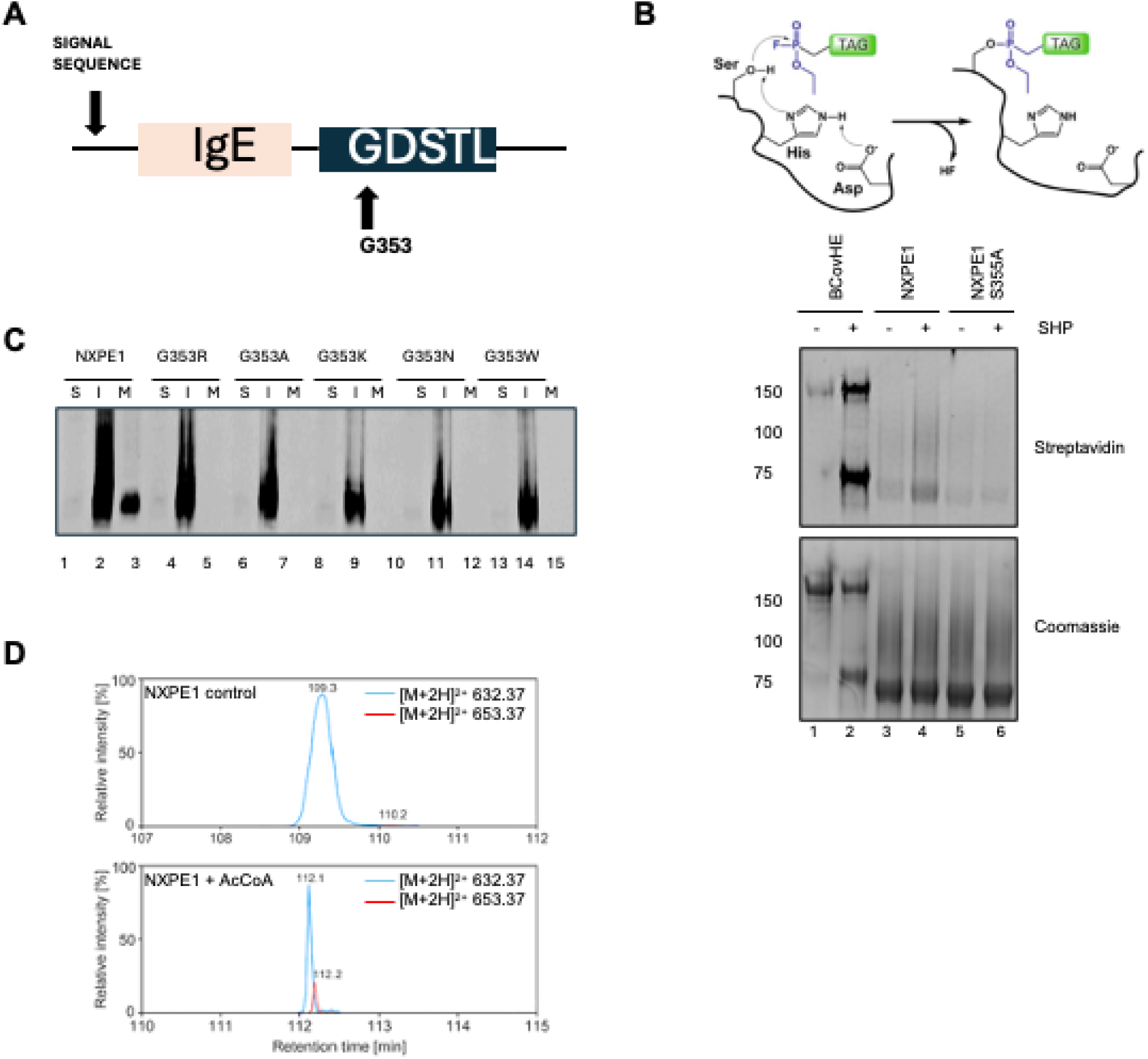
NXPE1 is an *O*-acetyltransferase and member of the serine hydrolase family. (**A**) Domain structure of NXPE1. Sequence homology suggests that NXPE1 is a serine hydrolase of the GDSL/SGNH family with a characteristic GDSTL active site sequence. The UC-protective variant (G353R) is highlighted. S355 is the active site covalent catalyst. (**B**) (Top) Schematic of the fluorophosphonate ABPP probe and reaction with NXPE1. (Bottom) The well-known serine hydrolase BCovHE (bovine coronavirus esterase, positive control), NXPE1, and NXPE1 S355A were incubated with and without the serine hydrolase probe (SHP). Active-site labeling was then monitored using a streptavidin-based Western blot. Native NXPE1, but not the S355A mutant, can be captured with the SHP. (**C**) Assessing soluble protein expression with G353 variants. Only the endogenous protein is soluble and secreted into the medium (‘M’), whereas the G353R/A/K/N/W variants are insoluble (‘I’) and not observed in the soluble crude extract (‘S’) nor secreted into the medium. (**D**) Formation of a covalent acetyl-enzyme intermediate captured by MS. Purified NXPE1 and S355A mutant were incubated with or without acetyl-CoA, subjected to SDS-PAGE, trypsin treatment, and analysis by HPLC-ESI-MS. Acetyl-adduct is only observed with native NXPE1 in the presence of Ac-CoA.

Serine hydrolases are one of the largest enzyme superfamilies and are capable of diverse enzymatic activities in multiple metabolic processes. To characterize the function of NXPE1, we investigated acetyltransferase activity by testing whether it could form a covalent acetyl-enzyme intermediate upon incubation with acetyl-coenzyme A (CoA) in the absence of an acceptor substrate. Upon incubation with acetyl-CoA, followed by proteolytic digestion, and analysis of the peptide fragments via liquid chromatography-coupled electrospray ionization-mass spectrometry (LC-ESI-MS), we found a peptide containing the active site residue S355 in its modified, acetylated form, as indicated by a 42 Da mass shift (**Figure 2D**). The mutant NXPE1 S355A, however, was not acetylated upon incubation with acetyl-CoA. These results indicate that NXPE1 is a novel acetyltransferase and that S355 serves as the active site covalent catalyst.

### NXPE1 mediates 9-*O*-acetylation of Neu5Ac with high substrate specificity

Acetyltransferases transfer an acetyl group from a common donor, such as acetyl-CoA, to an acceptor substrate (38, 39). To identify the acceptor substrate of NXPE1, we conducted a screen with various monosaccharides, given the Golgi localization of the enzyme. We tested 18 different sugars as potential acetyl group acceptors, including 11 nucleotidyl sugars and seven free monosaccharides (**Figure 3A**). After incubating NXPE1 with acetyl-CoA and each acceptor molecule, the reaction was acid-quenched to preclude nonenzymatic acetyl group transfer, precipitated NXPE1 was removed by centrifugation, and the sugars were modified with 1,2-diamino-4,5-methylenedioxy-benzene (DMB) to facilitate their detection. The substrate/products were then analyzed using high-performance liquid chromatography-coupled quadrupole time-of-flight mass spectrometry (HPLC-qTOF-MS). The results showed acetyltransferase activity with several sugars, the strongest occurring with *N*-acetylneuraminic acid (Neu5Ac) and Cytidine-5′-monophosphate-*N*-acetylneuraminic acid (CMP-Neu5Ac). Neu5Ac, the predominant sialic acid in human and many mammalian cells, is essential for various cellular functions, while CMP-Neu5Ac serves as a critical intermediate in the biosynthesis of sialoglycoconjugates (40, 41). Additionally, approximately 10-fold weaker acetyltransferase activity was observed with UDP-Xylose, a nucleotide sugar involved in the biosynthesis of glycosaminoglycans. NXPE1 also showed ∼100-fold weaker, but detectable turnover with several other sugars, such as *N*-acetylgalactosamine (GalNac) and UPD-galactose (**Figure 3A**). Importantly, no reaction was detected at all with the NXPE1 S355A mutant that disrupts the catalytic serine.

**Figure 3.**
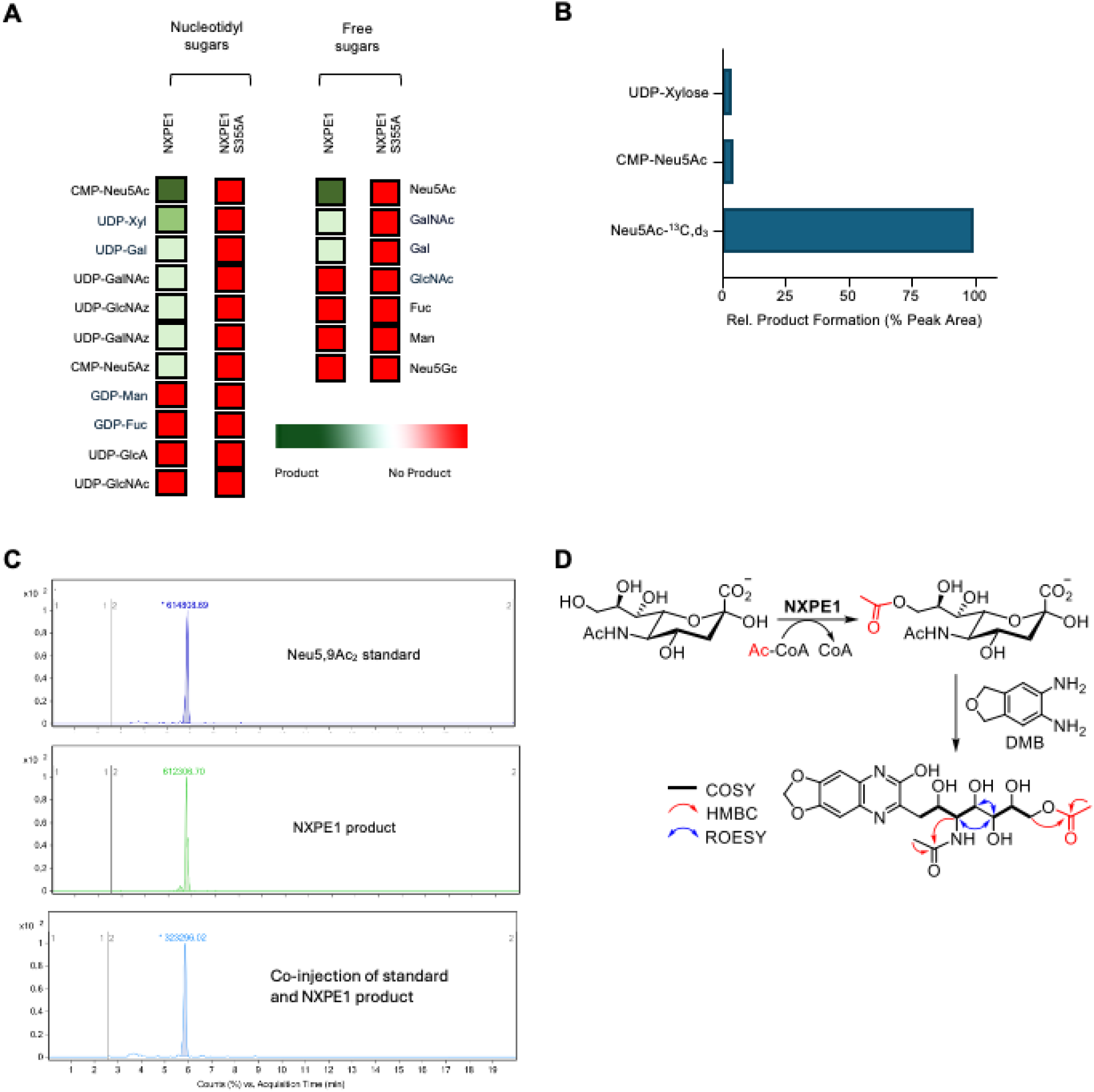
NXPE1 catalyzes 9-*O*-acetylation of Neu5Ac. (**A**) NXPE1 and the S355A mutant were screened against 18 acceptor substrates as indicated. The amount of acetylated product was then quantified using HPLC-qTOF-MS analysis, as indicated by the color bar. The data are normalized to the reaction of NXPE1 with Neu5Ac, which showed the highest turnover. (**B**) NXPE1 substrate specificity was determined by incubating it with equimolar concentrations of UDP-xylose, CMP-Neu5Ac, and Neu5Ac-^13^C,d_3_. The isotopically labeled form of Neu5Ac was used to distinguish it from CMP-Neu5Ac. Product formation was quantified using HPLC-qTOF-MS, which is shown in the bar graph and is normalized to Neu5Ac-^13^C,d_3_ (set to 100%). The average of three independent measurements are shown. (**C**) HPLC traces of DMB-labeled authentic Neu5,9Ac_2_ (top), the NXPE1 product (middle), and co-injection of the two samples. Coelution indicates the compounds are similar. (**D**) Reaction catalyzed by NXPE1 and analysis by DMB. NMR-based structure elucidation of the reaction product is shown with key NMR correlations highlighted.

To test the specificity of NXPE1, we incubated it with equimolar concentrations of the most active acceptors, leveraging isotopically labeled Neu5Ac (Neu5Ac-^13^C,d_3_) to distinguish it from CMP-Neu5Ac. Strikingly, NXPE1 showed high selectivity towards Neu5Ac, generating 20-fold higher levels of acetylated product than with any of the other substrates tested (**Figure 3B**). The activity was enzyme concentration-dependent with product accumulating in a time-dependent manner. No acetylated product was observed with NXPE1 S355A. Extended incubation of NXPE1 with Neu5Ac led to the appearance of small amounts of bisacetylated product (0.2–2% relative to monoacetylated Neu5Ac, see below).

Neu5Ac contains several alcohol functionalities that could be acetylated by NXPE1. To determine the precise location of acetylation, we conducted two experiments. First, we co-injected the product of the reaction with a standard consisting of Neu5,9Ac_2_, which was generated synthetically, and found the two compounds to co-elute, indicating they are identical (**Figure 3C**). Moreover, we carried out a large-scale reaction of NXPE1 with Neu5Ac, purified the product and determined its structure using 1D/2D NMR spectral analysis. The data were consistent with the analysis above and showed Neu5,9Ac_2_ as the reaction product. Specifically, a new acetyl group was detected in the product with δ_H_/δ_C_ of 2.05/20.2 ppm for the methyl group and δ_C_ of 174.2 ppm for the carbonyl-carbon. HMBC correlations from the methylene-protons at C-9 to the carbonyl carbon (δ_C_ 174.2) established the location of the modification (**Figure 3D**). Together, these findings show that NXPE1 is an acetyltransferase that acts preferentially on Neu5Ac and acetylates the C-9 hydroxy group to generate Neu5,9Ac_2_.

### NXPE1 mediates sialic acetylation of colonic mucus

Having established the biochemical activity of NXPE1, we next sought to assess its role in cell models using colonic organoids. Specifically, we developed organoids from five donors expressing the common allele of NXPE1 and seven donors homozygous for the UC-protective variant G353R. We also generated NXPE1^−/−^ organoids using CRISPR/Cas9 as a negative control. Since human cells also encode the paralogous NXPE4, which we reasoned may have redundant functions that could potentially compensate for NXPE1 KO, we also generated NXPE1^−/−^/NXPE4^−/−^ double KO organoids. The organoids were then assessed by western blot and/or MS analysis to determine the role of NXPE1. NXPE1 was only detected in organoids expressing the common allele but not those expressing the G353R variant, presumably because it is a hypomorphic allele encoding an unstable protein (**Figure 4A**).

**Figure 4.**
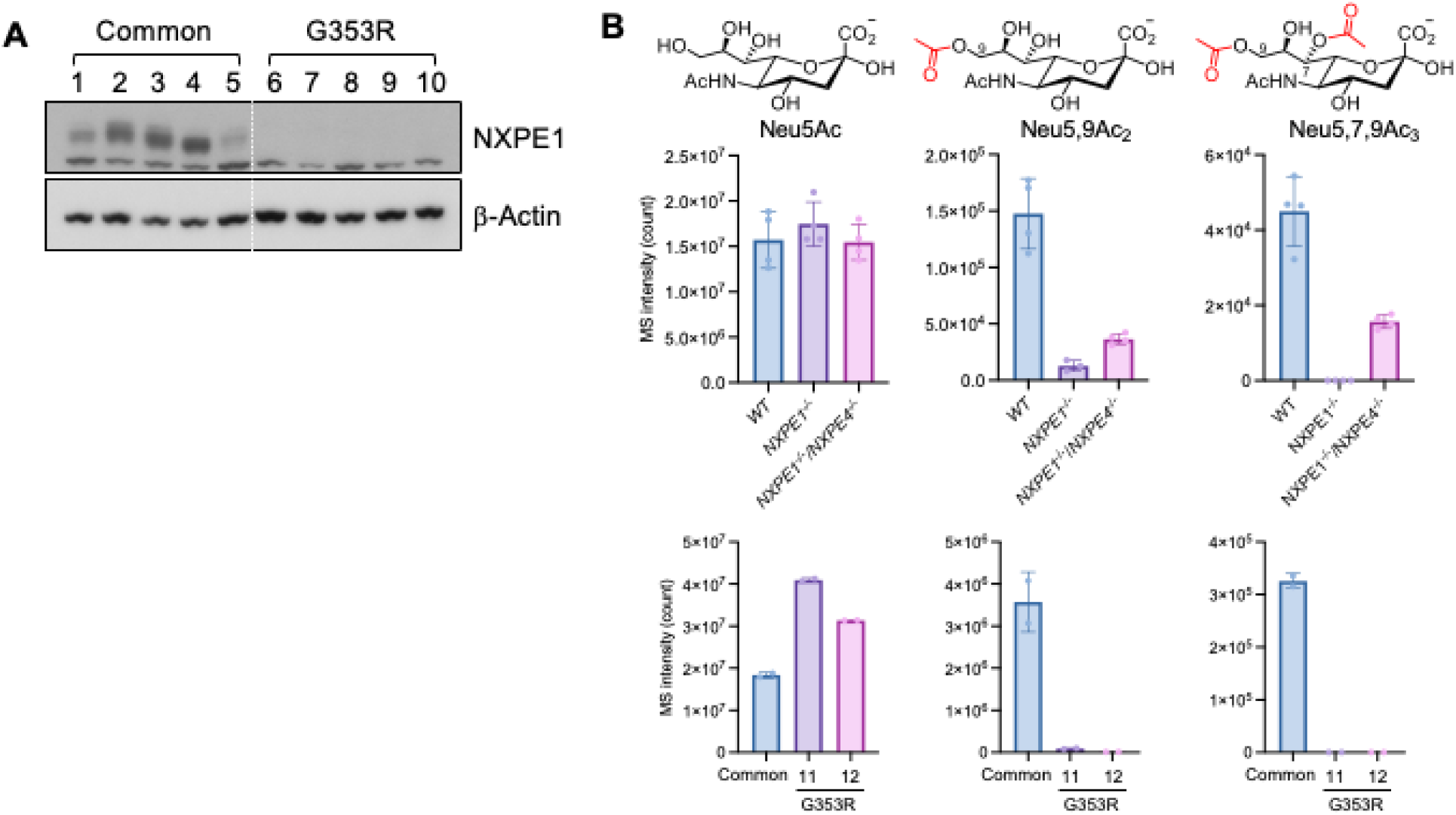
NXPE1 acetylates Neu5Ac in mucus from human colonic organoids, and G353R variant is a loss-of-function allele. (**A**) Immunoblot of NXPE1 using differentiated human intestine organoid lysates obtained from patients with the common allele of NXPE1 or UC-protective variant G353R. (**B**) Sialic acid content of organoids expressing endogenous NXPE1 or the G353R variant. Sialic acids were released from the outer layer secreted mucus and their distribution determined by HPLC-qTOF-MS. The sialic acid monitored is shown above each column. The average of five independent experiments is reported; error bars represent S.D.

To interrogate the sialic acid content of the organoids, we grew them as monolayers in air-liquid interface (ALI) cultures with vasoactive intestinal peptide (VIP), which promotes production of a hydrated apical mucus barrier. Accordingly, mucus was harvested and processed to release sialic acids, and we labeled these with DMB and assessed them by HPLC-qTOF-MS. Neu5Ac was detected in mucus from cells expressing the common allele, G353R variant, as well as the NXPE1^−/−^ and NXPE1^−/−^/NXPE4^−/−^ (**Figure 4B**). However, the product (Neu5,9Ac_2_) was produced at high levels only in mucus from cells expressing the common allele. It was undetectable in mucus from homozygous G353R variants and observed minimally in mucus from the NXPE1^−/−^ organoids. Importantly, mucus from NXPE1^−/−^/NXPE4^−/−^ cultures revealed a similar decrease in Neu5,9Ac_2_ as NXPE1^−/−^, although to a slightly lesser extent due to less efficient knockout of NXPE1 in double knockout organoids. Together these findings suggest that NXPE4 does not compensate for the loss of NXPE1, because they do not have redundant functions with respect to acetylation of sialic acid on mucus (**Figure 4B**). Furthermore, acetylation of sialic acids from NXPE1^−/−^ organoids could be rescued by incubating mucus from the knockout with recombinant NXPE1, further indicating that it is required and sufficient for this activity (data not shown). Detailed analysis of the HPLC-qTOF-MS data also revealed a second minor product, that we identified as a tris-acetylated neuraminic acid (Neu5,7,9Ac_3_); this product was also observed in biochemical assays of NXPE1 upon extended incubation with Neu5Ac and Ac-CoA (data not shown). Together, these results show that NXPE1 generates Neu5,9Ac_2_ and smaller amounts of Neu5,7,9Ac_3_, thus modulating the sialoglycome of colonic mucus. Given the recognized importance of mucus glycan modification in regulating the integrity of the intestinal barrier (42, 43), our findings offer new insights by identifying the UC-associated gene NXPE1 as a novel enzyme that is required for acetylation of sialic acids on mucins.

## DISCUSSION

We have discovered the reaction catalyzed by NXPE1, a protein with previously unknown function that was implicated by GWAS with a protective effect in UC patients. We find that NXPE1 is highly expressed in intestinal epithelial cells within the Golgi apparatus and that it is a serine hydrolase that can be captured via ABPP. NXPE1 catalyzes self-acetylation upon incubation with acetyl-CoA, a reaction that is diagnostic of an acetyltransferase. A substrate screen showed that NXPE1 transfers the acetyl group to the prevalent sialic acid Neu5Ac to generate the 9-*O*-acetyl product and, to a lesser extent, the 7,9-*O*-bisacetyl product. This reaction can modulate mucus viscosity in colonic organoids, a function that is abrogated in cells carrying the G353R variant, which we report is a loss-of-function allele. Interestingly, CASD1 and NXPE1 show opposing substrate preferences with NXPE1 selectively acetylating Neu5Ac in competition with CMP-Neu5Ac, whereas CASD1 catalyzes minimal acetylation of the free sugar with respect to the nucleotidyl CMP-Neu5Ac form. These acetyltransferases, therefore, exhibit disparate substrate preferences though similar localization in the Golgi.

The reaction catalyzed by NXPE1 and its loss-of-function variant are key to explaining its possible role in UC, a disorder that is associated with alterations of intestinal barrier function and dysbiosis of commensal flora. Mucus is produced in various body parts, and cells modulate mucus properties using several strategies. Our results herein suggest that acetylation of alcohol functionalities on sialic acids, which are subsequently appended onto mucins, is one such strategy that may modulate viscosity by altering mucus hydration and physical transitions between liquid and gel-like states. We propose that the UC-protective variant loses this ability and could thus produce a constitutively viscous mucus. In the context of UC, this can be beneficial as it produces a more rigid barrier and keeps bacteria that contribute to inflammation at bay. To gain further insights into this process and further assess this model, understanding the functions of other NXPE paralogs will be critical.

The corollary of the proposed model is that inhibition of NXPE1 may be beneficial in UC, and the enzyme therefore provides a new, attractive target that directly addresses intestinal barrier disruption. Therapeutic interventions to restore barrier integrity represent a significant unmet need and offer opportunities for the development of first-in-class drugs. Following our learnings from genetics and the mechanistic basis for the protective effects of NXPE1 G353R, we hypothesize that small molecule inhibitors of NXPE1 may mimic this protective effect on barrier function. The spatially and temporally well-defined expression profiles of NXPE1 suggest that inhibition would not have broad adverse off-target effects, though this would need to be tested. Aside from pointing to a possible target for UC treatment, our results more broadly highlight the power of deorphanizing human enzymes to uncover new biological processes that may be targeted therapeutically.

## METHODS

### General Experimental Procedures

An Agilent Technologies 1200 series HPLC instrument coupled 6120 quadrupole MS equipped with an electrospray ionization (ESI) source was used to obtain low-resolution HPLC-ESI-MS data. NMR spectra to determine the structure of the NXPE1 product were collected at the Princeton University Department of Chemistry NMR Core facilities. 1D/2D NMR spectra were recorded on a Bruker Avance 800 MHz NMR spectrometer carrying a multipurpose cryoprobe. Semi-preparative HPLC purifications for NMR experiments were performed on an Agilent 1290 Infinity Series HPLC system which was equipped with an automatic liquid sampler, a diode array detector, and an automated fraction collector.

### Homology modelling

The three-dimensional structure of NXPE1 was predicted using Alpha Fold. To improve its stability, we strategically truncated a sequence at T67 at the N-terminal region with low folding confidence. Subsequently, we introduced a secretion signal from CrypA to enhance the folding and secretion properties of this NXPE1.

### Expression and purification of NXPE1

Plasmids enabling the secreted expression of NXPE1 in Expi293 mammalian cells were based on a modified pRUSHmam vector (Invitrogen) carrying sequence stretches encoding an *N*-terminal signal peptide (CrypA) and a *C*-terminal TEV-TSFLAG tag. The sequence encoding residues E68–C547 of human NXPE1 was amplified by PCR. The resulting PCR products were ligated into the restriction sites of the modified pRUSHmam vector, resulting in the plasmids pRUSHmam-CrypA-T67-NXPE1-TEV-TS-FLAG, pRUSHmam-CrypA-T67-S355A-NXPE1-TEV-TS-FLAG, and pRUSHmam-Crypa-T67-G353RNXPE1-TEV-TS-FLAG. Expi293F human cells were grown at 37°C in shaking cultures at 125 RPM in Expi293™ Expression Medium (Thermo, A1435101) and maintained at a density of 0.8–4 x 10^6^ viable cells per ml.

Transient expression of WT NXPE1 and S355A-NXPE1 was performed using the ExpiFectamine™ 293 Transfection Kit system (Thermo Fisher), as described above. Conditioned medium was collected 96 h after infection, concentrated by centrifugation (5000g, 10 min, 4°C). The supernatant was then supplemented with BioLock and incubated for 30 min. It was then dialyzed against 50 mM sodium phosphate, 200 mM NaCl (pH 7.5) and passed over a Strep-Trap HP column (IBA). After washing with 20 column volumes of the same sodium phosphate buffer supplemented with 5% glycerol, bound protein was eluted by a desthiobiotin gradient. The desthiobiotin was subsequently removed by gel filtration using a Superdex 200 10/300 GL column equilibrated with 20 mM Tris-HCl, 200 mM NaCl, 5% glycerol, and 1 mM TCEP (pH 7.5).

### Differential Scanning Calorimetry (DSF)

All proteins (BCovHE, CASD1, WT NXPE1, and S355A-NXPE1) were used at a final concentration of 3 µM for this assay. SYPRO Orange (Invitrogen S6651) was used at a final concentration of 10X. Experiments were carried out in 20 mM Tris-HCl, 200 mM NaCl, 5% glycerol, 1 mM TCEP (pH 7.4). All DSF experiments were performed using an Eppendorf Realplex2 Mastercycler. Each sample was divided into four 10 µL replicates. Sample solutions were dispensed into 96-well optical reaction plates (Thermo Fisher Scientific 4306737), which were sealed with an optical PCR plate sheet (Thermo Fisher Scientific AB-1170). Fluorescence intensity was measured via the JOE emission filter (550 nm) and “PTS clear plate” was set as the background for the calibration. Temperature was continuously increased at a rate of 1°C/min. Melting curves were directly exported from the instrument and analyzed using Prism 6 (GraphPad Software Inc.).

### In vitro serine hydrolase assay

Serine hydrolase activity was measured in a total volume of 50 µl containing in 20 mM Tris-HCl, 200 mM NaCl, 5% glycerol, 1 mM TCEP (pH 7.5). The reaction contained 10 µM protein (BCoV-HE, WT NXPE1 or S355A-NXPE1) and 1 mM ActiveX Desthiobiotin serine hydrolase probe (SHP, Sigma #88317). The samples were incubated overnight at 37°C.

### In vitro formation of the acetyl-enzyme intermediate

Purified WT NXPE1 or S355A-NXPE1-S355A (5 µg) was incubated for 15 min at 37°C with or without 2.5 mM acetyl-CoA (Sigma Aldrich) in a 50 µl reaction in Tris buffer (20 mM Tris-HCl, 200 mM NaCl, 5% glycerol, 1 mM TCEP, pH 7.5). After storage at 20°C for at least 18 h to stabilize the acetyl-enzyme intermediate, the reaction mixture was separated by SDS–PAGE, the protein band was excised, digested with trypsin, and analyzed by HPLC-ESI-MS.

### In vitro sugar screen

Enzyme assays were performed for 3 h at 37°C in 60 µl containing 50 mM MES, 10 mM MnCl_2_, 0.5 mM acetyl-CoA (pH 6.5) with or without 3.8 µM WT NXPE1 or S355A-NXPE1. The following sugars were tested as acceptors at a concentration of 0.5 mM: CMP-Neu5Ac, GDP-L-fucose, GDP-D-mannose, GDP-xylose, UDP-galactose, UDP-glucoronic acid, UDP-galuronic acid, UDP-*N*-acetylglucosamine, UDP-*N*-acetylgalactosamine, UDP-*N*-acetylazidoglucosamine, UDP-*N*-acetylazidogalactosamine, CMP-*N*-acetylazidoneuraminic acid, 9-azido-Neu5Ac, CMP-*N*-glycolylneuraminic acid, Neu5Ac, galactose, *N*-Acetyl-D-glucosamine, fucose, mannose, N-glycolylneuraminic acid (Chemily Glycoscience). After 3 h, the reaction was quenched by addition of 12 µl of 2 M propionic acid and incubation at 80°C for 1 h. Propionic acid was used to reduce *O*-acetyl group loss/migration during this step. After 1 h, the quenched reaction was cooled to RT for 10 min, spun down to remove precipitated protein, and the sugars then labeled with DMB by addition of 9 µl of an 80 mM DMB stock (prepared in acetone), 5 µl of β-mercaptoethanol, and 5 µl of sodium dithionite (0.3 M stock prepared in water). The mixture was incubated for 2.5 h at 50°C. It was then spun down, filtered, and analyzed by HPLC-qTOF-MS using an analytical Phenomenex Polar C18 column (4.6 x 150 mm, 1.6 μm, 100Å) and a mobile phase consisting of water/acetonitrile supplemented with 0.1% (v/v) formic acid. Elution was carried out isocratically at 16% acetonitrile in water for 12 min at a flow rate of 0.15 ml/min. The amount of acylated product was determined by relative quantification via extraction of the desired product ion and integration of the area underneath the peak. The amount of product observed in the reaction of WT NXPE1 with Neu5Ac was set to 100%. The average of at least three independent experiments are reported for the substrate screen and other enzymatic assays.

### Substrate specificity experiment

To characterize the preferred substrate of NXPE1, a competition experiment was carried out where the exact same reaction conditions were used as described above, except that NXPE1 was incubated with equimolar concentrations of three substrates that gave the highest turnover in the substrate screen. These were Neu5Ac, CMP-Neu5Ac, and UDP-xylose. Because DMB treatment results in the same product from Neu5Ac and CMP-Neu5Ac, isotopically labeled Neu5Ac (Neu5Ac-^13^C,d_3_) was used to differentiate it from CMP-Neu5Ac. The reaction was worked up and the amount of product determined as described above in the substrate screen.

### Purification of DMB-Neu5,9Ac_2_

To determine the position of acetylation by NXPE1, large scale (8 ml) experiments were carried using the same conditions as describe above in the substrate screen. After formation of the DMB adduct, the sample was lyophilized to dryness, resuspended in 400 μl of 50% MeOH/Water and then subjected to semi-preparative HPLC purification using a Kinetex C18 column (Phenomenex, 5 μm, 10 x 250 mm). DMB-Neu5,9Ac_2_ eluted at a retention time of 26 min using an 80 min gradient at 2 ml/min starting with 5% acetonitrile in water and ending at 45% acetonitrile in water. The product was lyophilized to dryness and then subjected to 1D/2D NMR analysis.

### Human colonic organoid culture

Endoscopic biopsy samples were obtained from healthy donors enrolled in the Prospective Registry in IBD Study at MGH (PRISM). Written informed consent was obtained from study participants and the study protocols were reviewed and approved by the Mass General Brigham Human Research Committee (#2004-P-001067). Human colon spheroids were established and maintained as previously described (44) with a slight modification. Briefly, biopsy samples were washed with PBS containing penicillin (100 units/ml), streptomycin (0.1 mg/ml) and gentamicin (50 ug/ml) for 5 min at room temperature. After three washings with PBS, tissues were incubated with 5 mM EDTA in PBS for 45 mins at 4°C. After wash with PBS, several pipetting yielded supernatants enriched in colonic crypts. Crypts were pelleted by centrifugation at 200 g for 5 mins and then plated in 15 ul of Matrigel (Corning) and maintained in 500 ul of 50% L-WRN conditioned media supplemented with 10 uM Y-27632 (R&D systems, cat# 1254) and 10 uM SB431542 (Tocris Bioscience, 1614). Medium was changed every 2-3 days and organoids were passaged to 1: 3 using TrypLE every 7 days.

## ACKNOWLEDGMENTS

This work was supported by the National Institutes of Health (R.J.X., M.R.S.) as well as the Crohn’s and Colitis Foundation, the Center for Microbiome Informatics and Therapeutics, and the Klarman Cell Observatory (R.J.X.). This work was further supported by a GSK-Broad Institute research collaboration (R.J.X., D.B.G.). We thank members of the GSK team, including Nadia Eckert, Sunil Nagpal, Aaron Cheng, Karen Simpson, Tharini Selvakumar and Silvia Gonzalez-Del Valle for their scientific discussions regarding this work.

## Notes

### Competing Interest Statement

The authors have declared no competing interest.

